# Reproductive traits of the European catfish (*Silurus glanis*) during the early stages of invasion in the Lower Tagus River

**DOI:** 10.1101/2023.10.26.563989

**Authors:** Christos Gkenas, Diogo Ribeiro, João Gago, Diogo Dias, Chandani R. Verma, Pradeep Kumkar, Filipe Ribeiro

**Affiliations:** MARE, Centro de Ciências do Mar e do Ambiente / ARNET, Rede de Investigação Aquática, Faculdade de Ciências, Universidade de Lisboa, 1749-016 Lisboa, Portugal; Escola Superior Agrária, Instituto Politécnico de Santarém, Quinta do Galinheiro - S. Pedro, 2001-904 Santarém, Portugal; Department of Zoology and Fisheries, Faculty of Agrobiology, Food and Natural Resources, Czech University of Life Sciences Prague, 16500 Prague, Czech Republic

**Author notes:** Corresponding author: Christos Gkenas; ORCID ID 0000-0002-9863-2289; Diogo Ribeiro: ORCID ID 0000-0002-1820-3557 João Gago: ORCID ID 0000-0003-3893-5920 Diogo Dias: ORCID ID 0000-0003-2638-8923 Chandani Verma: ORCID ID 0000-0002-8577-8887 Pradeep Kumkar: ORCID ID 0000-0002-5443-1899 Filipe Ribeiro: ORCID ID 0000-0003-3531-5072.

**Keywords:** invasive, size at maturity, gonadosomatic index, fecundity, oocyte diameter, Siluridae

## Abstract

Freshwater ecosystems face severe challenges from biological invasions, leading to biodiversity loss, disruption of ecosystem services, and economic impacts. Human-mediated activities, such as aquarium trade and sport angling, contribute to species introductions, with potential negative consequences for native ecosystems. The European catfish (*Silurus glanis*) is one of the world’s largest freshwater fish and has been intentionally introduced into diverse regions, impacting native ecosystems. However, limited research exists on its reproductive traits outside its native range. This study addresses this gap by examining the reproductive characteristics of non-native European catfish populations in the Lower Tagus River in Portugal, focusing on size at maturity, spawning period, and fecundity. The observed balanced sex ratio aligns with studies of native populations. Variations in size at first maturity (TL_50_) among populations highlight the influence of habitat conditions, temperature, food availability, growth rate, and geographical location on this trait. The extended spawning season (March to June) in the Tagus River is consistent with native populations, but variations may occur based on environmental conditions and water temperature. Absolute fecundity ranged from 8,961 to 335,500 oocytes, showing positive relationships with body size and emphasizing the reproductive potential of European catfish in Portugal. Egg size variations, along with asynchronous egg development, contribute to the species’ reproductive strategy, favoring its invasive success. Management efforts should include monitoring, regulations on introductions, removal programs, and public awareness to mitigate their impact. Future research should focus on understanding how non-native European catfish populations adapt in various regions and continue to impact ecosystems.

## Introduction

Freshwater environments are among the ecosystems most impacted by biological invasions (Reid et al., 2019), experiencing an enormous loss of biodiversity (Bellard et al., 2016), while disrupting the supply of ecosystem services (Walsh et al., 2016) and causing substantial economic losses (Haubrock et al., 2022). Human-mediated activities drive biological invasions through multiple pathways of species introduction that operate at different spatial and geographic scales, including aquarium fish commerce, inland fisheries, maritime activity, aquaculture trade, and ornamental purposes (Tricarico, 2012; Carpio et al., 2019). However, fish intentionally introduced for economic or recreational purposes are released in locations where they are expected to thrive, as targeted introduction pathways prioritize species with desirable biological traits that favor their invasion success (Ruesink, 2005). For example, in sports fisheries introductions, a specific subset of species ranging from Salmonids and Cyprinids to top predator fishes exhibits biological traits that maximize their success (Ribeiro et al., 2008; Carpio et al., 2019). The practice of introducing non-native species is expected to continue, particularly in developed countries where sport angling serves as a significant additional pathway. Nevertheless, careful management is necessary to prevent further harm to ecosystems (Rahel & Smith, 2018; Carpio et al. 2019).

Life history traits, such as longevity, physiological tolerance, and early reproduction, significantly influence a species’ success in a new environment (Winemiller, 2005; García-Berthou, 2007; Ribeiro et al., 2008). However, the success of species invasions is strongly shaped by geographical factors and depends on key biological characteristics (Marr et al., 2010). For example, having a small body size, early maturation, high parental care, and shorter lifespan can contribute to the invasion success in the case of fish invasions in the Iberian Peninsula (Ribeiro et al., 2008; Alcaraz et al., 2005). Conversely, in different geographical contexts, invasive fish species with contrasting traits from native fish, such as high reproductive lifespans, exhibit greater invasive potential (Olden et al. 2006). Specifically, fish with larger body sizes, high fecundity, and long spawning seasons have a greater capacity to produce a larger number of oocytes throughout their lifetime, thereby increasing their likelihood of successful establishment and subsequent spread (Wootton and Smith, 2015; Liu et al., 2017).

The European catfish, *Silurus glanis*, is ranked among the 20 largest freshwater fish worldwide, with a maximum length of 2.8 meters and a weight of 120 kg (Boulêtreau & Santoul, 2016). It exhibits rapid growth, reaching a length of up to 1 meter by the age of six and can live for up to 70 years in the wild (Copp et al., 2009; Bergström et al., 2022). Due to its large size, it has become a popular target for recreational anglers, resulting in deliberate introductions in Western and Southern European countries (Boulêtreau & Santoul, 2016; Vejřík et al., 2019; Castagné et al., 2023), as well as in other countries such as China, Tunisia, and Brazil (Cucherousset et al., 2018). The species is commonly found in reservoirs and higher-order rivers near submerged objects (Gago et al., 2016) and exhibits tolerance to variations in water temperature, low levels of dissolved oxygen, high pollutant concentrations, and salinities of up to 15 ppm (Carrasco et al. 2011; Huertas et al. 2016). Furthermore, *S. glanis* has a wider niche and an opportunistic diet compared to other predatory fish (Vejřík et al., 2017), allowing it to colonize new ecosystems. It adapts to new resources and its diet can vary across seasons and during different stages of the invasion (de Santis & Volta, 2021). The impacts of invasive catfish populations on the ecosystem can be significant, as they engage in competition and predation with native species, alter nutrient cycling, and affect overall ecosystem health (Cucherousset et al., 2018; Vejřík et al., 2017).

The European catfish has been the subject of extensive research due to its ecological and economic impact as a top predator in freshwater systems, as well as its value as a recreational fishing resource (see Copp et al., 2009; Cucherousset et al., 2017; Vejřík et al., 2019). However, despite the large number of studies on the species’ predatory behaviour, there is a lack of research on its reproductive traits outside its native region. Hence, the current study aims to examine whether the reproductive characteristics of non-native *S. glanis* could explain its invasion success in a Mediterranean country, namely describing the size at first maturity, fecundity, and spawning period of the species. Additionally, this study compares these findings with existing information on the reproductive biology of the European catfish in its native range and other invaded regions to comprehend the ecological factors contributing to successful invasions. Conducting such research can provide valuable insights for invasive risk assessment and the development of effective management policies.

## Materials and methods

Sampling campaigns were conducted over a span of six consecutive years (2017-2023) in the Lower Tagus River, Portugal (Lat: 39°4’19.17″N-39°40’3.32″N; Long: 8°45’38.12″W-7°30’52.16″W). Monthly samples of *S. glanis* were collected using electrofishing, gill nets, and baited hook-lines, or were obtained from professional fishermen’s catches. Fish sampling of European catfish did not occur in December due to a combination of unfavorable winter conditions that posed significant challenges and safety concerns for conducting sampling activities. We pooled the data based on a preliminary analysis that revealed the absence of statistically significant differences among different years and months.

Upon collection, the catfish were immediately placed on ice and subsequently frozen for later laboratory examination. In the laboratory, individual catfish specimens were defrosted and measured for total length (TL, to 0.1 cm) and weighed for total weight (TW, with a precision of 0.01 g) and eviscerated weight (EW, to 0.01 g). Gonads were removed to determine the sex and subsequently weighted (GW, to 0.001 g).

The sex ratio was calculated monthly and compared to the hypothetical 1:1 proportion using a chi-square (χ^2^) test (with Yates’ correction). We estimated female size at maturity at which 50% of the females are mature (TL_50_). A logistic regression analysis was used by fitting the total length as an explanatory variable and the stage of maturity (0□=□immature, 1□=□mature) as the binomial response variable. Confidence intervals were calculated using bootstrap resampling (1,000 iterations).

Gonad maturity phases were determined by examining gonad development at the macroscopic scale, following the criteria outlined by Alp et al. (2004). The six macroscopic maturity stages were (I) immature, i.e., sex not distinguishable with the naked eye; (II) not ripe; (III) maturing; (IV) ripe; (V) running ripe; and (VI) post-spawning/spent. The gonadosomatic index (GSI) was calculated by dividing the wet weight of the gonads by the total wet weight of the fish: GSI = 100 × (G/TW), where G is the weight of the gonads (0.1 mg) and TW is the fish total weight (0.01 g). To evaluate monthly variations in the GSI of females, we applied the Kruskal-Wallis test as the data did not follow a normal distribution.

Fecundity was determined based on 60 mature-ripe females (stages IV and V) using the standard gravimetric method. We obtained three sub-samples from the front, middle and caudal sections of each gonad and counted the number of oocytes in each sub-sample to estimate the absolute fecundity (AF) using the formula: AF = (Gonad weight x Egg number in the subsample/Gonad subsample weight) (Alp et al. 2004). Relative fecundity (RF) was calculated from the formula RF = AF/TW, by dividing the absolute fecundity with the total weight of fish. Regression analyses were used to describe the relationship of fecundity to the fish total length and body mass. To determine the oocyte diameter, we measured the diameter of 90 oocytes in ripened gonads during the breeding season using a stereoscope and Image J, an image analysis program. Examining oocyte diameter during the reproductive period will enable us to assess the duration of the reproductive season and determine whether this species exhibits multiple spawning or single spawning behaviour. Variation in fecundity and oocyte diameter among months was analysed with analysis of covariance (ANCOVA) using fish length as the covariate. In order to meet normality and homoscedasticity assumptions of absolute fecundity and oocyte diameter, the data were previously log_10_-transformed. All data processing and analysis were performed using R version 4.2.2 (R Core Team, 2022).

A comprehensive literature review was also conducted to investigate the reproductive biology of European catfish in both their native and nonnative habitats. This review primarily focused on elucidating the relationships among key life history traits, including but not limited to size at maturity, fecundity, reproductive season, and their respective correlations within the specified geographic regions. Furthermore, the potential influence of environmental factors, such as temperature and latitude, on these reproductive traits was also discussed within the framework of this study.

## Results

In total, we assessed 819 *S. glanis* specimens ranging from 10.5 to 210 cm (TL) of which 238 females (32.4 - 163 cm), 201 males (34.3 - 210 cm) and 380 immature (10.5 - 98 cm) - whose sex was not recognized macroscopically. The overall sex ratio (females/males = 1.2:1) did not significantly deviate from the expected 1:1 ratio (χ^2^ = 3.118, p = 0.077), and this pattern remained consistent throughout the observed months (Supplementary Table S1). The smallest mature female measured 57 cm TL and the estimated size at 50% sexual maturity (TL_50%_) was 79.2 cm TL (95% CI = 77.1 – 81.3 cm TL) using binomial regression (Figure 1).

**Figure 1:**
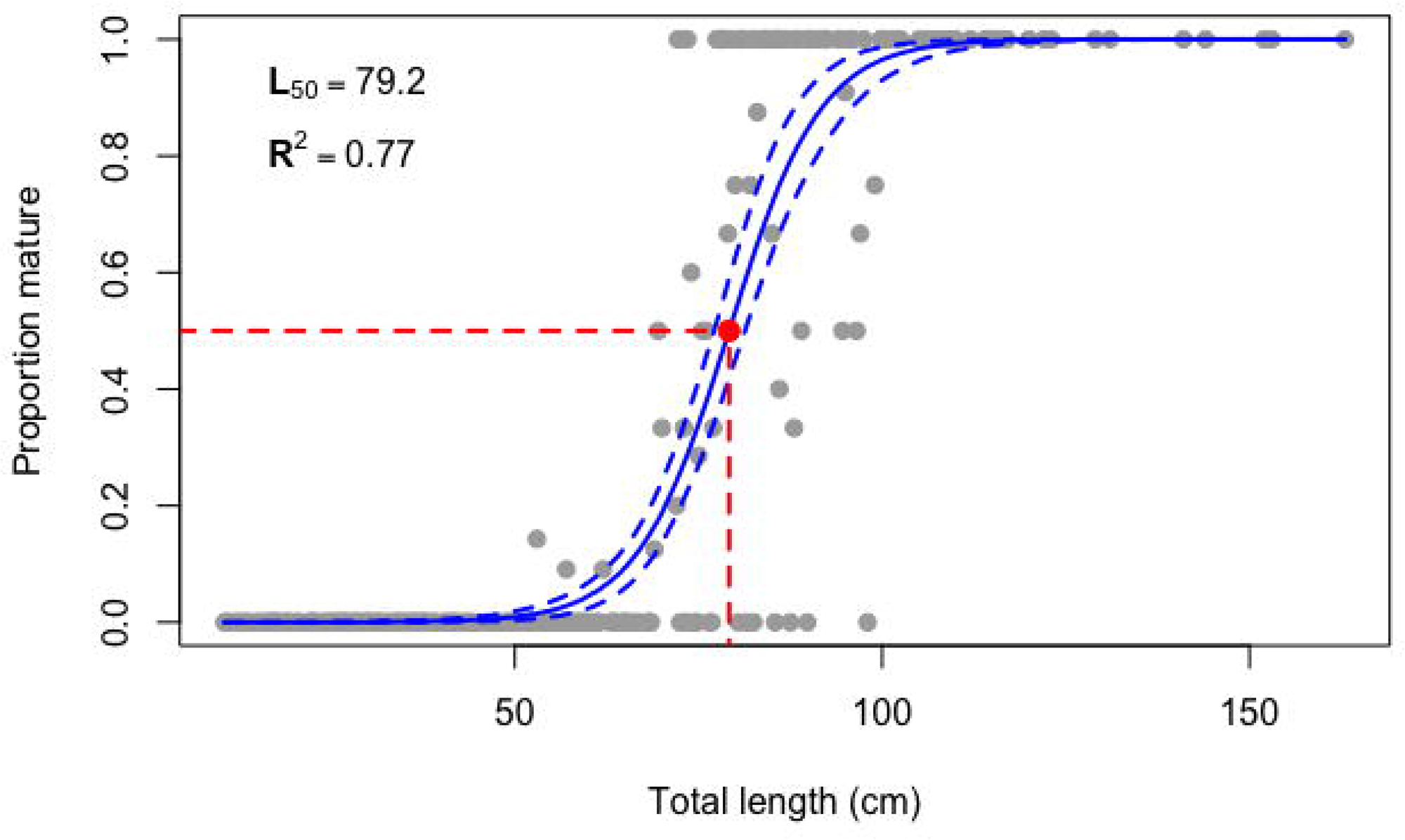
Logistic regression curve of the proportion of mature female *Silurus glanis* with total length in the Lower Tagus River. The solid line represents the model-predicted values, and the dashed lines represent the corresponding 95% confidence intervals and red lines indicate respective values on the figure.

The GSI of female European catfish displayed significant variation along months (Kruskal Wallis H, χ^2^ = 81.96, p < 0.001), spanning a range from 0.1% to 17.4%. Starting in January, there was a notable increase in mean GSI values, which reached its peak in March (8.9%) followed by a gradual decline to 1.2% by November (Figure 2). These observations indicate that the spawning period for female *S. glanis* extended from April to June, possibly commencing in March, as indicated by the highest GSI values observed. However, there were some variations in the GSI values in September and October, suggesting potential reproductive activity by some females.

**Figure 2:**
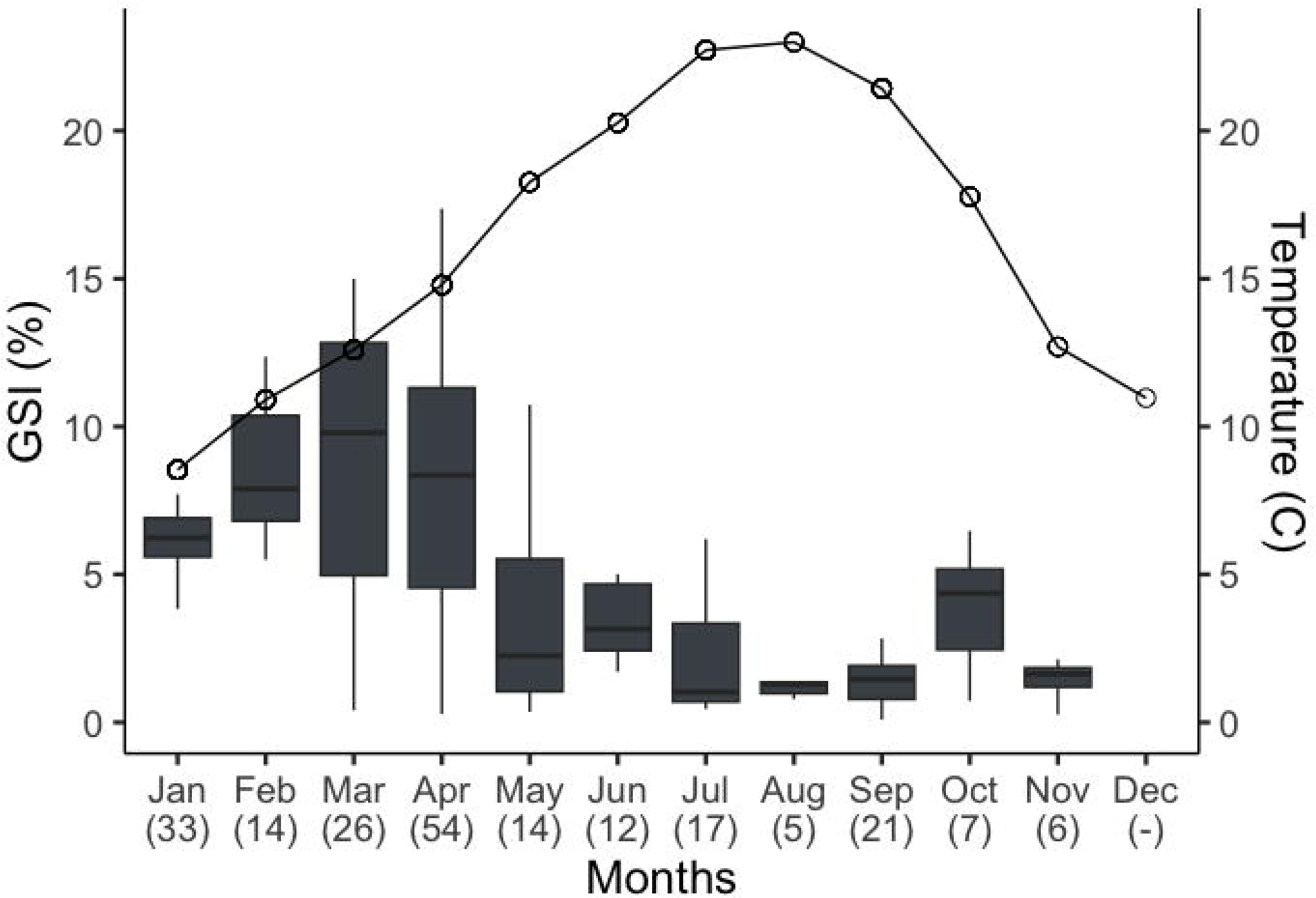
Box plots illustrating monthly variations in gonadosomatic index (GSI %) and temperature (^0^C) for female *Silurus glanis* from the Lower Tagus River. The box represents the interquartile range (IQR; 25th and 75th percentiles), and the line within the box is the median. Whiskers represent the 75th percentile + 1.5 x IQR and the 25th percentile + 1.5 x IQR. Temperature values are shown as mean, extracted from the SNIRH database (available at https://snirh.apambiente.pt/).

During the breeding period, a total of 60 ovaries at stage IV and stage V were collected to calculate the fecundity. Absolute fecundity exhibited a wide range, varying from 8,961 to 335,500 oocytes per female, with a mean of 78,657 (±SD 63,515). Relative fecundity showed variation from 6.2 to 28.5 oocytes per gram of TW, averaging 14.2 (±SD 4.7). Absolute fecundity increased significantly with TL (ANCOVA, F_5,54_ = 59.508, P < 0.001) and TW (ANCOVA, F_5,54_ = 64.611, P < 0.001), as supported by high R^2^ values observed in the regression analysis between absolute fecundity and TL as well as TW (Figures 3a and 3b). No significant relationships were found between relative fecundity and TL (P > 0.05, R^2^ = 0.01) or TW (P > 0.05, R^2^ = 0.01).

**Figure 3:**
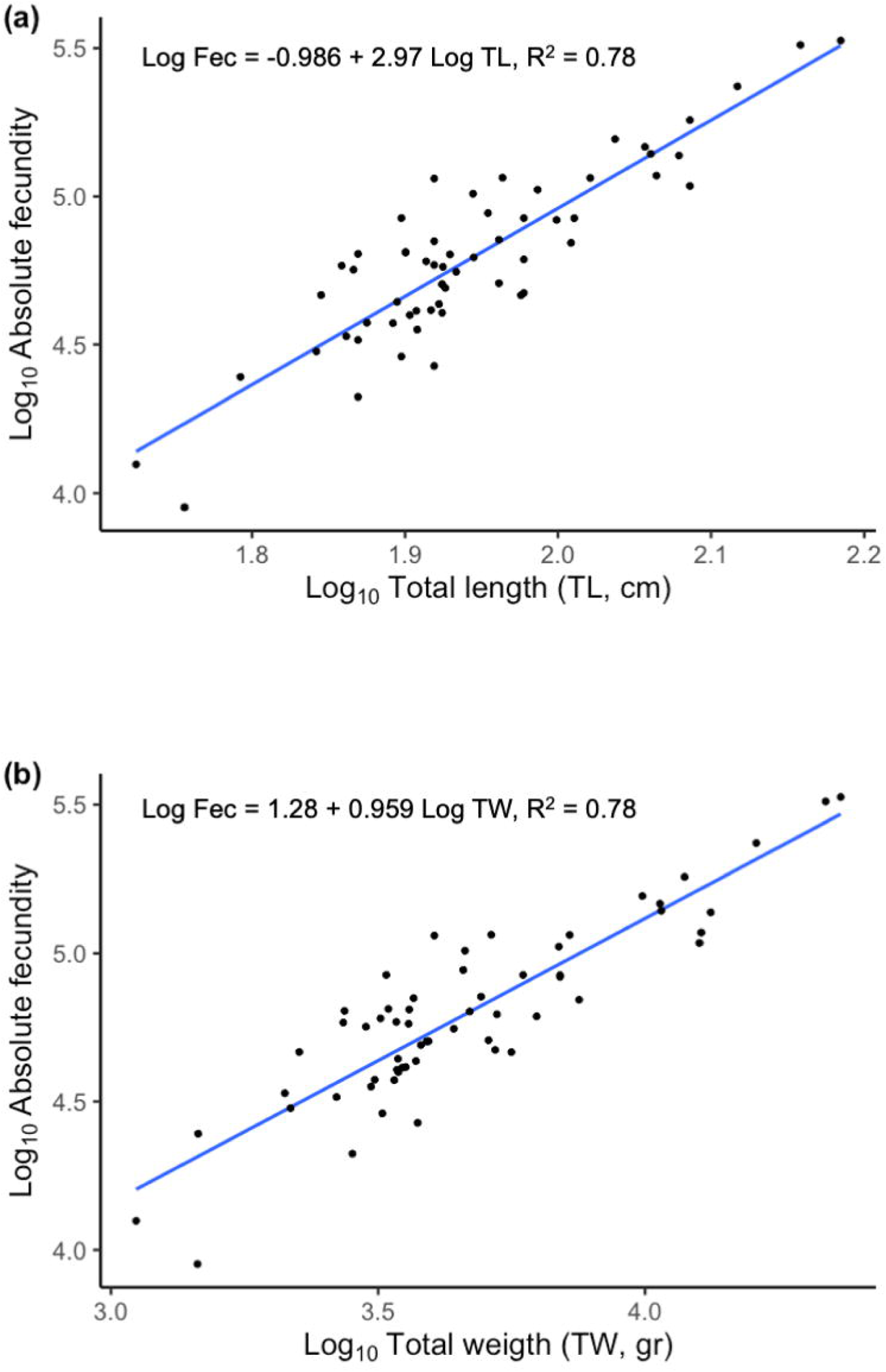
Relationships between absolute fecundity and (a) total length (TL, cm) or (b) total weight (TW, gr) of female *Silurus glanis* in the Lower Tagus River.

The mean diameter of oocytes ranged from 0.13 to 3.21 mm (mean = 1.94 ±SD 0.40), and did not show significant differences among the anterior, middle, and posterior sections of the ovary (χ^2^ = 2.77, P > 0.05). Oocyte size-frequency distributions for oocyte diameters were continuous and without interruption, indicating asynchronous development and suggesting more than one spawning event during the reproductive period (Figure 4). Notably, the modal classes of larger oocytes (>2.25 mm) seemed to appear in different months for each female size class, with smaller females exhibiting earlier occurrences in March-April, and larger females in April and May (Figure 4). The oocyte diameter - fish length relationship varied significantly across months (F_5,5913_ = 20.923, P < 0.001). The maximum average size was observed in April (2.1 ±SD 0.4 mm).

**Figure 4:**
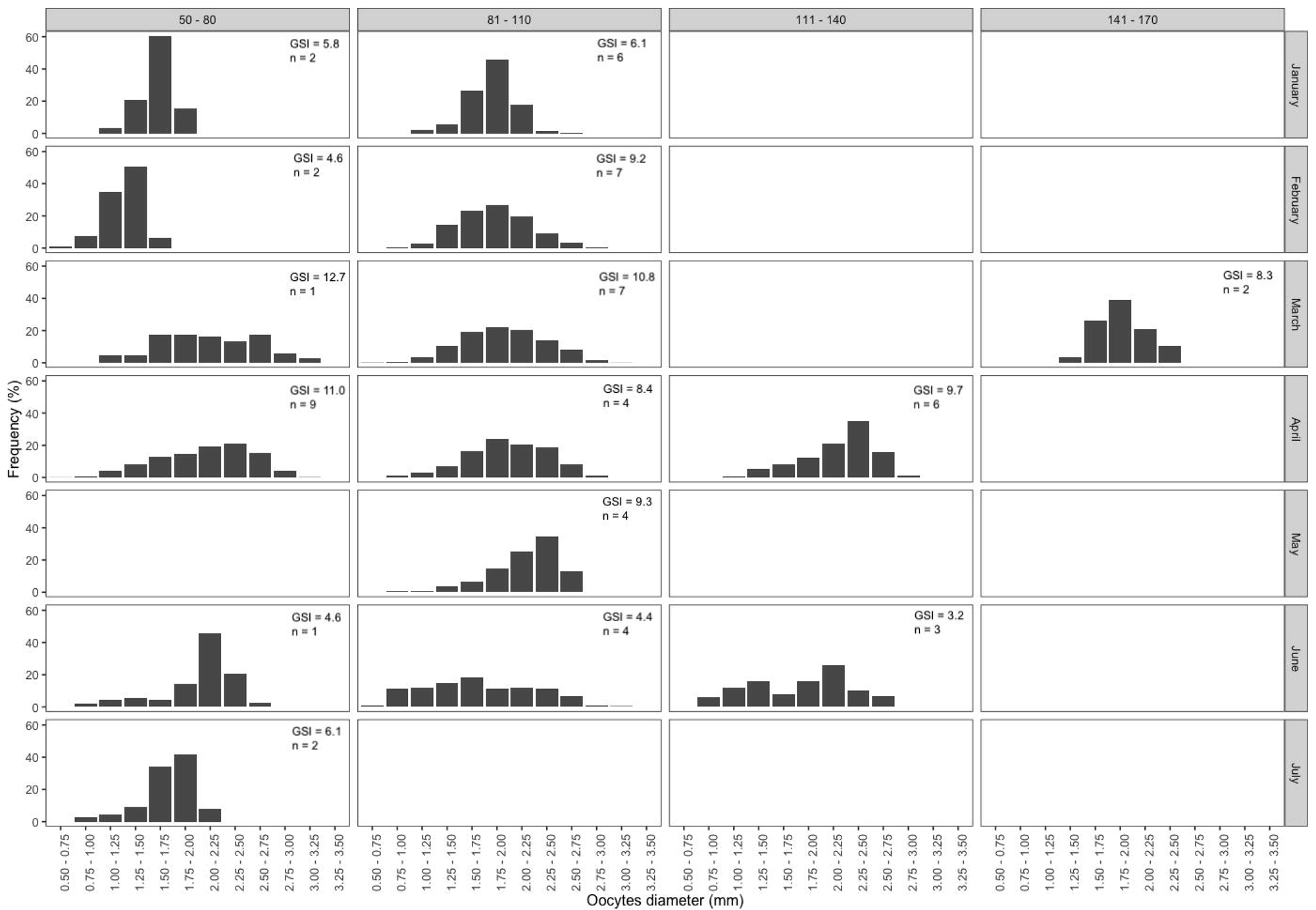
Monthly size frequency distributions of oocytes from female *Silurus glanis* at the Tagus Lower River, per size class; n: corresponds to the number of females analyzed for oocyte diameter, and GSI: represents the mean Gonadosomatic Index of the analyzed females.

## Discussion

The European catfish exhibited several biological traits, such as large size and extended longevity, that could have limited its establishment in Iberian freshwaters, while other traits, such as opportunistic predatory behaviour and parental care associated with high fecundity, could have facilitated its invasion of Iberian watersheds (Ribeiro et al., 2008; Alcaraz et al., 2005). Reproductive biology studies of the European catfish were primarily conducted on native populations, with most research conducted during the last century (Table 1). The current study contributes significantly to our understanding of the catfish reproductive biology, being one of the first studies conducted in its invaded range (but see Puzzi et al., 2003), and presenting an extensive sampling (Table 1). The European catfish population of the Tagus River exhibited an extended spawning season (March to June), early maturity (TL_50_ = 79.2 cm), high reproductive investment (GSI max = 17.4%), associated with a large number of oocytes produced (from 8,961 to 335,500 oocytes), and large oocyte diameters.

**Table 1:**
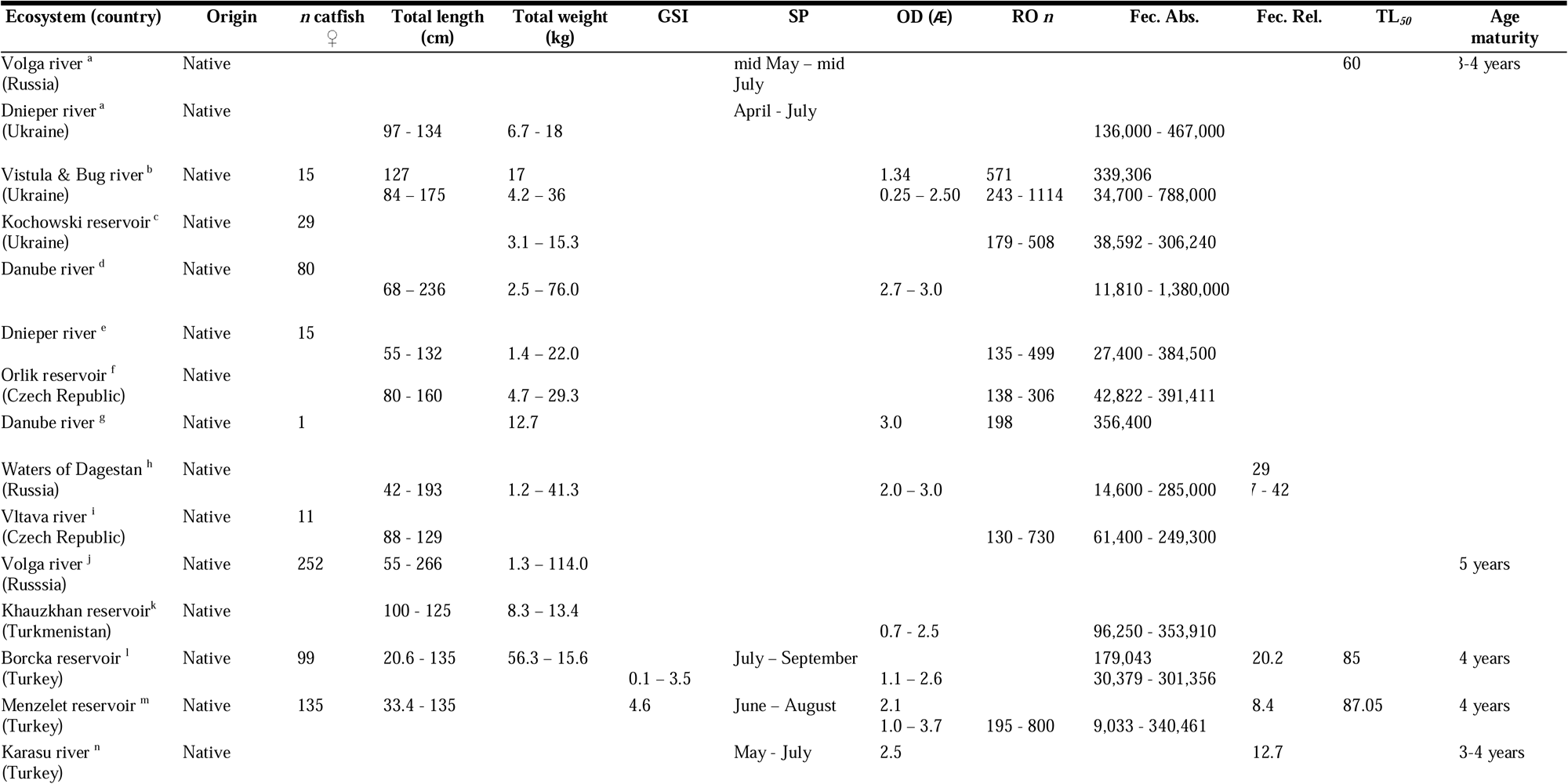
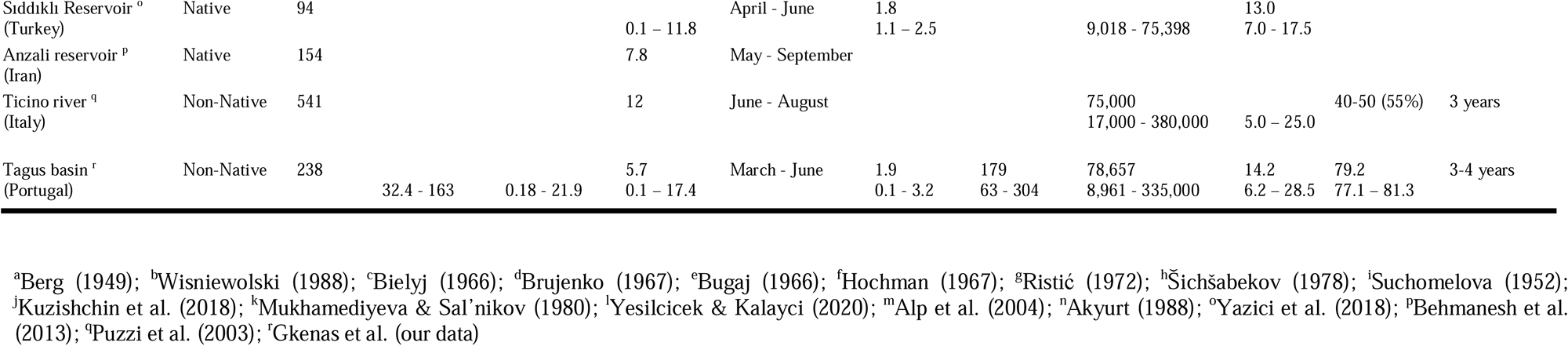
Summary of reproductive traits of European catfish (*Silurus glanis*) obtained from a literature search of native and non-native populations, with reference to Country, Drainage, Habitat type (Delta, River, Reservoir), Origin (Native, Non-Native), n - number of individuals analysed, TL - Range of Total length (cm), TW - Range of Total Weight (g), GSI - Gonadosomatic Index in %, SP - Spawning period observed in months, OD - Mean Oocyte Diameter in mm; ROn - Relative Oocyte number per gram; Fec. abs - Absolute fecundity (nr. of oocytes);Fec. Rel. - Relative Fecundity (nr. of oocytes/Kg of female weight), TL50 - Total Length at maturity of 50% of females (cm), with respective reference number in footnote;

This study involved a substantial number of analysed females (N=238), being one of the largest studies on European catfish reproductive biology (Table 1). However, during late fall and early winter, we were unable to collect fish samples due to severe winter conditions. Although Copp et al. (2009) suggested that this period corresponds to the gonad resting phase, our results indicate that catfish gonads are maturing, observed by the GSI values in January (Figure 2). Therefore, a comprehensive evaluation of the maturation cycle throughout the year is needed, particularly because in its invaded range in the Iberian Peninsula, catfish populations are subjected to higher water temperatures during the winter (9-11°C), compared to their native range (4-6°C) (Figure 2). Nevertheless, our consistent and frequent samplings of European catfish over the years enable a more robust assessment of their reproductive traits, addressing potential temporal variations. Finally, our findings highlight the importance of comprehensive size-at-maturity data, even in the absence of histological information. This observation aligns with previous studies (Alp et al., 2004; Yazici et al., 2018) and provides valuable insights into catfish biology. However, future research may consider incorporating histological data to further enhance our understanding of European catfish reproductive biology.

The sex ratio recorded in the current study was found to be balanced, which is consistent with the findings of previous reproductive studies of *S. glanis* within its native range (Alp et al., 2004; Yazici et al., 2018). Nevertheless, Yesilcicek & Kalayci (2020) reported a bias in the sex ratio, deviating from the expected 1:1 pattern. These sex-related differences have been attributed to several factors, including varying habitat characteristics, sampling period, and the use of different fishing gears and methods, which may selectively remove individuals from the larger size classes, thereby biasing sex ratios.

In our study area, we demonstrated a size at first maturity (TL_50_) of 79.2 cm for females, which, according to the growth curves reported by Alp et al. (2004) in the Menzelet reservoir, corresponds to fish that are approximately 4 years old. In Western Asia, several authors (Hochman, 1967; Abdullayev et al., 1978; Šichšabekov, 1978; Mukhamediyeva & Sal’nikov, 1980; Orlova, 1989) reported a TL_50_ range of 39 to 71 cm, corresponding to fish aging between 3-4 years. However, in the Volga Delta, *S. glanis* began maturing at age 2 (mean TL = 50.7 cm), with almost all fish becoming mature at age 6, and mass maturation occurring at ages 3-4, corresponding to 57-66 cm TL (Orlova, 1989). These findings highlight significant differences in the size at first maturity for females in the Lower Tagus River, being higher compared to other Western Asian populations but smaller than populations in the Menzelet Reservoir (Alp et al., 2004) and the Borçka Dam (Yesilcicek & Kalayci, 2020) in the East region of Turkey, where females matured later at 87.05 cm TL and 85.0 cm TL, respectively (Table 1). Indeed, variations in the TL_50_ among European catfish populations could be attributed to various factors, such as habitat conditions, temperature, food availability, growth rate, and geographical location, all significantly influencing the size at maturity of *S. glanis* (Copp et al., 2009; Yesilcicek & Kalayci, 2020). Furthermore, a previous study in the Po River in Italy showed some variation in the size at maturity among European catfish populations in non-native ranges, with 55% of females reaching maturity at smaller sizes ranging between 40 and 50 cm TL (Puzzi et al., 2003). Consequently, these observations of early maturation in European catfish populations in non-native habitats may represent an adaptive response, potentially enhancing their invasive success.

The reproductive period of the European catfish in the Lower Tagus River, spanned from March to June, as evidenced by the monthly gonadosomatic index (GSI) changes, corresponding to water temperatures ranging from 13°C in March to 18°C in May and reaching 20°C in June. This finding is consistent with native *S. glanis* populations reproducing between April and June in the Sıddıklı Reservoir (Yazici et al., 2018) and from May to June in Karasu stream, in Turkey (Akyurt, 1988). However, extended timing has been noted from June to August in native catfish at Menzelet Reservoir (Alp et al., 2004) and from July to August in nonnative catfish in the Po River (Puzzi et al., 2003). In colder regimes, in Eastern Europe, spawning occurred from April to July in the Dnieper Delta and mid May to mid July in the Volga Delta (Berg, 1949), coinciding with 18–22 °C water temperatures (Copp et al., 2009). During this research, we observed that a prolonged temperature increase in August and September coincided with an extension of the reproductive activity among certain female individuals in the Lower Tagus River. This observation supports the findings of Maitland and Campbell (1992) and Greenhalg (1999), which suggest that European catfish spawning can be extended due to the effects of temperature and day length. Emphasizing the significance of ecological conditions and water temperature regimes in determining the spawning season, it is suggested that the timing and the duration of reproduction may vary depending on latitude, habitat characteristics and water temperature.

Absolute fecundity increased with increasing length and mass, following an expected pattern observed in fishes (Winemiller & Rose, 1992). Similar positive relationships between fecundity and total length or weight have been documented in European catfish populations from both native (Akyurt, 1988; Alp et al., 2004; Yazici et al., 2018; Yesilcicek & Kalayci, 2020) and non-native areas (Puzzi et al., 2003). In our study, *S. glanis* exhibited high fecundity, with absolute fecundity ranging from 8,961 to 335,500 oocytes, which is within the range of estimated fecundities reported in previous studies (Puzzi et al., 2003; Alp et al., 2004; Yazici et al., 2018; Yesilcicek & Kalayci, 2020; see Table 1). However, our absolute fecundity estimates were lower than those reported from the Vistula (Wisniewolski, 1988) and Danube rivers (Brujenko, 1967), being two to three times higher, respectively. Thus, all data indicated a variability in estimates within and between studies (Table 1). We recorded a mean estimated relative fecundity of 14.2 oocytes per gram (6.2 – 28.5 oocytes per gram). Similar estimates have been observed for catfish in the Menzelet Reservoir (mean 8.4 oocytes per gram) (Alp et al., 2004), the Karasu River (mean 12.7 oocytes per gram) (Akyurt, 1988), the Siddikkli Reservoir (13 oocytes per gram) (Yazici et al., 2018), and the Borcka Reservoir (mean 20.2 oocytes per gram) (Yesilcicek & Kalayci, 2020). However, catfish from Eastern Europe had the highest recorded estimates at a mean of 29 oocytes per gram (7–42 oocytes per gram) (Šichšabekov, 1978) (see Table 1). Therefore, the relative fecundity of European catfish varied significantly, influenced by factors such as food availability, water temperature, fish length, and geographic location.

In this study, we observed that the oocyte size of *S. glanis* ranged from 0.13 mm to 3.21 mm, with an average oocyte diameter of 1.94 mm. Furthermore, our analysis revealed a significant positive relationship between oocyte size and both fish length and months, indicating that larger females tend to produce larger oocytes. This observed pattern in oocyte size aligns with findings from populations of European catfish in Eastern Europe (Wisniewolski, 1988; Brujenko, 1967; Bugaj, 1966; Šichšabekov, 1978; Mukhamediyeva & Sal’nikov, 1980) and the Eastern region of Turkey (Alp et al., 2004; Akyurt, 1988; Yazici et al., 2018) (Table 1). Moreover, the size frequency distributions of oocytes revealed that females retained the ability to spawn due to the asynchronous development of oocytes. Notably, mature females were found to contain substantial remnants of small oocytes during spawning. This observation suggests that a significant proportion of oocytes may not complete full maturation and are not released but rather regress within the fish (Puzzi et al., 2003). This asynchronous development of oocytes is a common reproductive strategy in freshwater catfish, allowing females to spawn multiple batches of oocytes over time, ensuring that at least some oocytes will survive and hatch even in unpredictable environments, and adjusting the size and number of oocytes they produce in response to environmental conditions.

In summary, this paper represents one of the few studies on the reproductive traits of the European catfish outside of its native range. It provides new insights into the reproductive dynamics of *S. glanis* in their non-native habitat, showing an extended reproductive period and possibly an early maturation. The observed values of fecundity and size at maturity contribute to our understanding of this invasive fish potential adaptive strategy considering the broad variation of the several reproductive indicators (Table 1). In its invaded range, urgent population control actions are needed and these should incorporate this biological knowledge. In addition to continuous monitoring and evaluation of catfish populations, it will be necessary to observe changes in reproductive traits and behaviour over time, given that non-native fishes could change their biological traits in response to changes in population density (Bohn et al., 2004). Implementing catfish population control measures should involve fishing techniques that reduce bycatch (Vejřík et al., 2019) and target catfish aggregation sites (Cucherousset et al., 2012). These actions aim to increase the efficiency of control measures and mitigate their ecological impact. In this regard, population control strategies should primarily target adult and juvenile catfish sizes, which can be achieved using fishing techniques (e.g. long-lines or large mesh size nets) designed to catch individuals larger than 50 cm, focusing on capturing adult individuals or late juvenile individuals. Future research should encourage international collaboration among managers to enable coordinated catfish removal actions in shared rivers, such as the Tagus River. Establishing long-term monitoring programs, combined with dedicated local population control efforts, is essential for monitoring the progression and impact of European catfish. This is particularly important, as observed in the lower Po River (Castaldelli et al., 2013) where a long term monitoring program described the extirpation of two native fishes several decades after catfish invasion. Furthermore, considering Tagus endemisms like the *Iberochondrostoma olisiponense* (Veríssimo et al., 2018), these extinctions could have global implications, especially in endemic-rich drainages that are highly invaded by predatory fishes, such as the Lower Tagus River (Ribeiro et al., 2021; Magalhães et al., 2023).

## Author contributions

CG: conceptualization, data analysis and visualization, writing - original draft; DR: methodology, laboratory process, writing – review and editing; JG: methodology, laboratory process, writing – review and editing; DD: methodology, laboratory process, writing – review and editing; CV: methodology, laboratory process, writing – review and editing; PK: methodology, laboratory process, writing – review and editing; FR: investigation, conceptualization, resources; writing – review and editing; All authors read and approved the final manuscript.

## Acknowledgements

The authors would like to thank all the fishermen who contributed samples, which made this study possible. Special thanks go to Carlos Serras, João Lobo, and Francisco Pinto. The authors are also thankful for the assistance provided by several volunteer students who helped in fish processing over the years, namely Beatriz Castro, Mafalda Moncada, and Sofia Nogueira.

## Funding information

This study was conducted within the framework of the projects SONICINVADERS (FCT ref. PTDC/CTA-AMB/28782/2017) and MEGAPREDATOR (FCT ref. PTDC/ ASP-PES/4181/2021), co-funded by the European Commission under the EU LIFE Nature & Biodiversity 422 Project programme (Project 101074458 — LIFE21-NAT-IT-PREDATOR). Additional support was provided by Fundação para a Ciência e a Tecnologia (FCT) through the projects UIDB/04292/2020 and UIDP/04292/2020 awarded to MARE and through project LA/P/0069/2020 granted to the Associate Laboratory ARNET. C. Gkenas (DL57/2016/CP1479/CT0036) and F. Ribeiro (CEEC/0482/2020) are supported by individuals contracts from FCT.

## Conflict of interest statement

The authors declare no conflicts of interest.

## Data availability statement

The data presented in this study are available on request from the corresponding author.

## Notes

### Competing Interest Statement

The authors have declared no competing interest.

